# Genetic transformation of the dinoflagellate chloroplast

**DOI:** 10.1101/452144

**Authors:** Isabel C. Nimmo, Adrian C. Barbrook, Jit Ern Chen, Katrin Geisler, Alison G. Smith, Manuel Aranda, Purton Saul, Ross F. Waller, R. Ellen R. Nisbet, Christopher J. Howe

## Abstract

Coral reefs are some of the most important and ecologically diverse marine environments. At the base of the reef ecosystem are dinoflagellate algae, living in symbiosis with coral. Efforts to understand the relationship between alga and coral have been greatly hampered by the lack of an appropriate dinoflagellate genetic transformation technology. By making use of the plasmid-like fragmented chloroplast genome, we have introduced novel genetic material into the dinoflagellate chloroplast genome. We have shown that the introduced genes are expressed and confer the expected phenotypes. Genetically modified cultures have been grown for five months with subculturing, maintaining the introduced genes and phenotypes. This indicates that cells continue to divide after transformation and that the transformation is stable.

## Introduction

Coral reefs are complex ecosystems, made up of many thousands of species. At the base of the ecosystem are dinoflagellate algae, frequently referred to as zooxanthellae. These single-celled algae live in symbiosis with corals as intracellular photosynthetic symbionts, providing fixed carbon to the host. Loss of the symbiotic alga results in coral bleaching, which is one of the most urgent and worsening worldwide ecological concerns. In 2016, 85% of the Great Barrier Reef was found to be affected by coral bleaching, a significantly higher proportion than had been previously identified ^1^.

Change in sea water temperature is recognised as one of the environmental causes of coral bleaching^2^. It is likely that this results in disturbance of photosynthetic electron transfer in the dinoflagellate symbiont and consequent damage^3,4^. The PsbA (D1) reaction centre protein of photosystem II is believed to be an important target of such damage^5^. The key subunits of photosynthetic electron transfer chain complexes, including the PsbA protein, are encoded in the dinoflagellate chloroplast genome^6^. There have been no reports to date of transformation of the dinoflagellate chloroplast genome, hampering attempts to study the mechanism of bleaching.

An alternative approach to transformation of the chloroplast might be to insert genes for proteins carrying chloroplast targeting sequences into the nucleus. There have been numerous attempts at stable nuclear transformation of dinoflagellates, but none has been clearly successful. An early report of the transformation of the dinoflagellates *Amphidinium sp.* and *Symbiodinium microadriaticum* mediated by silicon carbide whiskers, with selection for resistance to hygromycin or G418 and using β-glucuronidase (GUS) as a reporter, appeared to produce transformants after 12 weeks^7^. However, there are no reports of successful use of this technique since the initial publication. More recently, Ortiz-Matamoros and co-workers reported transient expression of GFP in *Symbiodinium* using a plasmid designed for plant transformation introduced by treatment with glass beads and polyethylene glycol, and selection for resistance to the herbicide Basta (gluphosinate)^8^. However, these cells were not capable of cell division, and no genetic confirmation of transformation was carried out. Transformation with the same plant plasmid mobilised by the plant pathogen *Agrobacterium* was also reported, although the transformed cells again failed to divide^9^. The lack of stable expression of heterologous genes limits the use of these techniques for functional biochemical studies.

Here, we describe a method for stable transformation of the dinoflagellate chloroplast. The chloroplast genome of dinoflagellate species containing the carotenoid peridinin (which is the largest group and includes those forming symbionts with coral) is typically fragmented, comprising approximately twenty plasmid-like DNA molecules of 2-5 kbp known as ‘minicircles’^10,11^ Each minicircle typically carries a single gene, together with a conserved core region which is assumed to contain the origin of replication as well as the transcriptional start site^6^. Each chloroplast contains multiple copies of each minicircle, although the exact number varies according to the growth stage of a culture^12^.

We exploited this unusual minicircular genome organisation to create shuttle vectors for dinoflagellate chloroplast transformation. We created two artificial minicircles, both based on the *psbA* minicircle from the dinoflagellate *Amphidinium carterae*,, as the *A. carterae* chloroplast genome is the best characterized amongst dinoflagellates^13-15^. We replaced the *psbA* gene with a selectable marker (either a modified version of *psbA* which confers tolerance to the herbicide atrazine^16^, or a gene for chloramphenicol acetyl transferase, which confers resistance to chloramphenicol), and an *E. coli* plasmid backbone (to allow propagation in *E. coli*) and introduced these artificial minicircles into dinoflagellates using particle bombardment. Following selection, we could detect the presence of the artificial minicircles, and transcripts from them, using PCR and RT-PCR. Cultures under selection continued to divide and maintain the artificial minicircles for three months, indicating that transformation was stable. The availability of a method for dinoflagellate chloroplast transformation enables a range of studies on the maintenance and expression of this remarkable genome and the proteins it encodes, such as PsbA.

## Methods

### Culturing of *Amphidinium carterae*

*A. carterae* CCMP1314 (from the Culture Collection of Marine Phytoplankton) was cultured in f/2 medium on a 16h light/8h cycle, 18°C at 30μE m^−2^ s^−1^, as described previously^17^.

### Design of artificial minicircles

The pAmpPSBA artificial minicircle (predicted to confer atrazine tolerance) was prepared by PCR amplification of the wild-type *psbA* mincircle with outward facing primers from a point immediately downstream of the proposed poly-U addition site (Genbank AJ250262, fwd primer 1128-1155, rev primer 1127-1106)^13^. The linear PCR product was purified and cloned into the pGEM-T Easy plasmid (Promega), which contains the ampicillin resistance marker and a
bacterial origin of replication. The point mutations necessary to confer atrazine tolerance were introduced in a further round of PCR with *Pfu* polymerase and the following mutagenic primers, forward primer GTCTTATCTTCCAGTATGCTGGCTTCAACAACTCCCGTTCTC, reverse primer GAGAACGGGAGTTGTTGAAGCCAGCATACTGGAAGATAAGAC. This altered a TCC (Serine) codon to a GGC (Glycine) codon at position 260 of the PsbA protein (numbered as in AJ250262). The PCR products were treated with DpnI to digest any template DNA and then used to transform chemically competent *E. coli* JM109. Ampicillin selection was used to identify colonies containing pAmpPSBA, and plasmids were sequenced. A plasmid map is shown in Figure 1A.

The pAmpChl vector was synthesised by GeneArt. This vector was based on a pMA vector backbone, and contained the psbA minicircle (as above), but with the *psbA* coding region removed and precisely replaced by an *A. carterae* chloroplast codon-optimized *E. coli* chloramphenicol acetyl transferase gene (CAT)^18^, Figure 1B.

**Figure 1.**
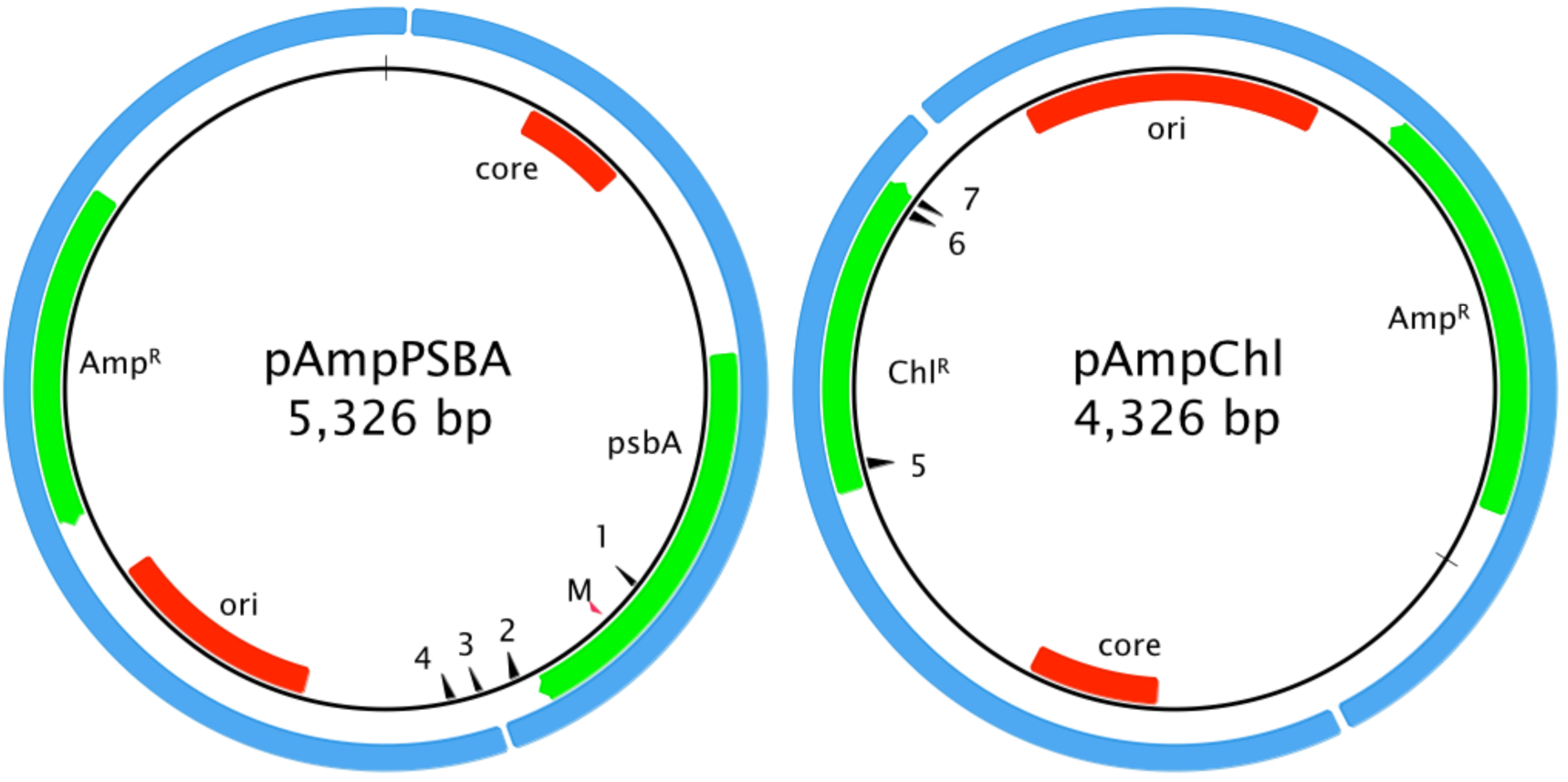
Artificial minicircle design. Left, pAmpPSBA. Right, pAmpChl. Origins
of replication (ori for *E. coli* plasmid, core region for *A. carterae* minicircle) are shown in red. Protein-coding genes (encoding ampicillin resistance, chloramphenicol resistance or PsbA) are shown in green. The blue shows the original source of of the genetic material (*E. coli* plasmid and *A. carterae* minicircle). The red arrow showing the position of the mutation in *psbA* that confers resistance to atrazine is marked with ‘M’. Primer sites pAmpPSBA 1: MCpG-F-II, 2: MC-pG-F, 3: MC-pG-R-II, 4: MC-pG-R; pAmpChl: 5: CAT-R, 6: CAT-F, 7: CAT-FSS. CAT-F-Nest and CAT-R-Nest are immediately adjacent to CAT-F and CAT-R, respectively.

Both vectors were propagated in *E. coli* under ampicillin selection, and isolated using the Promega Maxiprep plasmid purification protocol prior to transformation into *A. carterae*. The vectors were verified by DNA sequencing before use.

### Biolistic transformation of *A. carterae*

Biolistic transformation was carried out using a Biorad Biolistics PDS-1000/He system, Biorad rupture disks, stopping screens and macrocarriers. Preparation of particles carrying DNA was carried out using Seashell Technology’s DNAdel^TM^ gold carrier delivery system and 550 nm diameter gold particles.

*A. carterae* cells were grown to early log growth phase before harvesting prior to transformation. For each transformation, ∼2.5×10^7^ cells (as determined by light microscopy utilising a haemocytometer) were spotted onto the centre of a 1% agarose f/2 medium plate and allowed to dry. 0.5 mg of gold particles and 0.5 μg of vector DNA were used for each plate to be transformed. Each plate was shot using the above mentioned Biorad Biolistics PDS-1000/He system and rupture disks of either 1,100 PSI, 1,350 PSI or 1,550 pounds per square inch (p.s.i.) (see Table 1 for details).

Cells were immediately resuspended in 30-50 ml fresh f/2 medium and allowed to recover before the addition of the selective agent. Cells shot using the pAmpPSBA artificial minicircle were allowed 16-24 hours to recover. Cells shot using the pAmpChl artificial minicircle were allowed 72 hours to recover. Cells were maintained in liquid culture as they do not grow on solid medium. Media was changed every 4 weeks (atrazine) or 2 weeks (chloramphenicol). Cells were subcultured (1:2) every 8 weeks (atrazine) or 4 weeks (chloramphenicol). A step-by step protocol is described in ‘Biolistic Transformation of Amphidinium’ (https://www.protocols.io/view/biolistic-transformation-of-amphidniumhnmb5c6).

Culture survival was assessed by microscopy. A spot of 50μl was placed onto each of three microscope slides for each culture. After covering with standard coverslips, the entire volume for each was examined using a light microscope at x100 magnification. Cells were assessed as living if they showed more than simple Brownian motion. In general, ‘dead’ cells appeared to disintegrate shortly after movement ceased. If no living cells were found three days in a row, the culture was recorded as dead on day three.

### Extraction of DNA and RNA from *A. carterae*

Total RNA was isolated from *A. carterae* using the Trizol - chloroform method. Purification was carried out using the RNA clean-up with on-column DNase protocol of the Qiagen RNeasy kit as described by Rio, 2010^19^ except that isopropanol precipitation was carried out overnight, rather than for 10 minutes.

DNA was released from cells prior to PCR by resuspending 5 x 10^4^ to 10^7^ cells (depending on the number available) in 50 μl dH2O with ∼10-20 acid-washed 500μm glass beads and vortexing for 10 minutes.

### RT-PCR and PCR

First strand synthesis of the RNA was performed using Invitrogen Superscript IV using the manufacturer’s protocol and either random hexamer primers or a gene specific primer. Negative controls lacking reverse transcriptase were performed by the same method but replacing the reverse transcriptase enzyme with dH_2_O. PCR was carried out using Promega GoTaq polymerase according to the manufacturer’s instructions, and annealing temperature, extension time and MgCl_2_ concentration were varied as appropriate.

MC-pG-F GACTCTTAGAACGACTAGGCTTTCTG

MC-pG-R GCGTAATCATGGTCATAGCTGTTTC (Tm 55°C, 1 min extension time)

MC-pG-F-II GTATCCTCTTCTCTCCTTGC

MC-pG-R-II CTCTCCCATATGGTCGAC (Tm 50°C, 1.5 min extension time)

CAT-FSS ACCTTGCCACTCGTCAC

CAT-F GATATTTCTCAGTGGCATCGTAAG

CAT-R GCTCGTTAAGCATACGACCTAC (Tm 60°C,1.5 min extension time)

CAT-F-Nest CATCGTAAGGAACATTTCGAG

CAT-R-Nest CGACCTACATGGAAACCATC (Tm 50°C, 1.5 min extension time)

### Cloning and sequencing of PCR products

PCR products were separated by 1-1.5% agarose gel electrophoresis and visualized by staining with GelRed. PCR products were purified from excised gel pieces using the MinElute gel extraction kit (Qiagen). Some PCR products were directly sequenced after gel extraction whilst others were ligated into the pGEM-T Easy plasmid vector (Promega), following the manufacturer’s instructions. The ligation mix was used to transform chemically competent *Escherichia coli* TG1, followed by overnight growth on 1.5% LB agar containing ampicillin at 100 μg/ml. Individual colonies were picked and grown overnight in LB containing ampicillin at 100 μg/ml. Plasmids were extracted from resulting cultures using the QIAprep Spin Miniprep Kit (Qiagen). All sequencing was carried out using an Applied Biosystems 3130XL DNA Analyser in the Department of Biochemistry, University of Cambridge sequencing facility.

## Results

### Construction of artificial minicircles

Two artificial minicircles were used in this study. The first, pAmpPSBA, was designed to confer atrazine tolerance. Tolerance to atrazine in plants can be conferred by a single residue change in the PsbA protein, where a Serine is mutated to a Glycine^20^. We therefore cloned the *A. carterae psbA* minicircle into the *E. coli* vector pGEM T easy (Promega) and introduced the necessary mutations into the *psbA* gene using site-directed mutagenesis, Figure 1A. The second artificial minicircle, pAmpChl, was designed to confer chloramphenicol resistance. It is also based on the *A. carterae psbA* minicircle, but the *psbA* gene was excised and replaced by a *A. carterae* codon-optimized gene encoding chloramphenicol acetyl transferase. The plasmid backbone is E. coli pMA, Figure 1B.

### Biolistic transformation with pAmpPSBA

Nine experiments to introduce the pAmpPSBA artificial minicircle into *A. carterae* using biolistic transformation were carried out, using a range of rupture disk pressures. Each experiment was carried out in triplicate, and included a single negative control line (cells subjected to biolistic treatment but without pAmpPSBA). The mean survival time for each culture under selection was assessed by bright field microscopy, and results are shown in Table 1. In six experiments, *A. carterae* cells shot with particles carrying pAmpPSBA showed greater mean survival time under selection conditions than untransformed cells, suggesting successful transformation (Experiments A2, A3, A4, A5, A7 and A8). One experiment (A6) was harvested prior to the death of the control strain, so no conclusions can be drawn on relative survival times. Finally, two experiments showed no difference in the length of time for which cells survived. The first (Experiment A1) was carried out using the lowest pressure rupture disks. In this experiment, cells survived just 13 days, suggesting that cells had not been transformed, perhaps because an insufficient bombardment velocity had been applied. In the second experiment (A9), selection was carried out with 1 μg ml-1 atrazine, below the lethal level of 2 μg ml-1. Both test and control cultures survived at least three months, with subculturing occurring at eight week intervals.

**Table 1.** Biolistic transformation of *A. carterae* with pAmpPSBA. Each experiment was carried out in triplicate, thus producing three potentially transformed lines. In addition, one line of cells was subjected to biolistic bombardment, but without the pAmpPSBA (‘untransformed’). Note that cultures from experiments 5-9 were harvested for genetic analysis, and thus the listed survival time is the day of harvesting, labelled with *.

| Experiment | Rupture disk (p.s.i.) | Atrazine concentration ($\mu\text{g ml}^{-1}$) | Survival untransformed (days) | Mean survival pAmpPSBA (days) |
| --- | --- | --- | --- | --- |
| A1 | 1100 | 2.5 | 13 | 13 |
| A2 | 1350 | 2.5 | 13 | 16 |
| A3 | 1350 | 2.5 | 13 | 15 |
| A4 | 1550 | 2.5 | 13 | 17 |
| A5 | 1550 | 2.5 | 12 | 20* |
| A6 | 1550 | 2 | 7* | 7* |
| A7 | 1550 | 2 | 13 | 13* |
| A8 | 1550 | 2 | 12 | 15* |
| A9 | 1550 | 1 | 3 months* | 3 months* |

### Biolistic transformation with pAmpChl

Transformation attempts were also made with *A. carterae* using chloramphenicol resistance as selectable marker. Experiments were carried out with pAmpChl and 1550 p.s.i. rupture disks. In the first experiment, chloramphenicol (final concentration 10-50 μg ml-1) was applied after three days in liquid culture, to allow time for initial synthesis of chloramphenicol acetyl transferase (Experiments C1A-E), Table 2. No untransformed wild type cells (i.e. shot with gold particles without DNA) survived after 15 days, whatever the chloramphenicol concentration. However, at 10 μg ml-1 chloramphenicol, cells shot with particles carrying the pAmpChl plasmid survived for at least 35 days, Experiment C1A). When chloramphenicol concentration was 30 μg ml-1 or greater, cells shot with particles containing the pAmpChl plasmid had died by day 15, (Experiment C1C-E), Table 2. Note that where appropriate, cells were subcultured after 28 days.

**Table 2.** Biolistic transformation of *A. carterae* with pAmpChl. Each experiment
was carried out in triplicate, thus producing three potentially transformed lines. In addition, one line of cells was subjected to biolistic bombardment, but with gold particles lacking the pAmpChl (‘untransformed’). For experiment 1, cells from each plate (three shot with gold particles carrying the plasmid and one with gold particles only) were divided into five separate samples, each incubated at a different chloramphenicol concentration. Note that cultures from experiments C1A, C2, and C3, were harvested for genetic analysis, and thus the listed survival time of lines still alive at that point is the day of harvesting, labelled with *. Experiment C4 was still alive at 20 weeks and is thus marked with +.

| Experiment | Rupture disk (p.s.i.) | Chloramphenicol concentration ($\mu\text{g ml}^{-1}$) | Survival untransformed (days) | Mean survival pAmpChl (days) |
| --- | --- | --- | --- | --- |
| C1A | 1550 | 10 | 15 | 35* |
| C1B | 1550 | 20 | 15 | 17 |
| C1C | 1550 | 30 | 15 | 15 |
| C1D | 1550 | 40 | 15 | 15 |
| C1E | 1550 | 50 | 13 | 15 |
| C2 | 1550 | 10 | 13 | 13* |
| C3 | 1550 | 10 | 14* | 14* |
| C4 | 1550 | 20 | 16 | 20 weeks + |

### Detection of artificial minicircles using PCR

To test if the transformation construct could be recovered from putatively transformed cultures, DNA was isolated from atrazine-selected *A. carterae* cultures (experiment A5, two lines designated A5.1 and A5.2) by vortexing with glass beads. DNA was also isolated from wild type *A. carterae* as a negative control. In addition a DNA purification was carried out with transformed cells (line A5.1), but without vortexing with glass beads (‘unbroken cells’) in order to test whether DNA remained adsorbed to the outside of cells. A positive control was included using the pAmpPSBA artificial minicircle. PCR was performed using the primers MC-pG-F and MC-pG-R (Figure 1) which lie on either side of the junction between the *psbA* minicircle and the pGEM-T Easy vector. A single product was amplified from lines A5.1 and A5.2, with no product detected from either the wild type or the ‘unbroken cells’ (Figure 2A). This product matched the size of product from the positive control. Both products were cloned and sequenced. The sequence was as expected from pAmpPSBA, as a chimaera between the psbA minicircle and the pGEM-T Easy vector, confirming that the atrazine-resistant *A. carterae* did indeed contain the pAmpPSBA plasmid.

**Figure 2.**
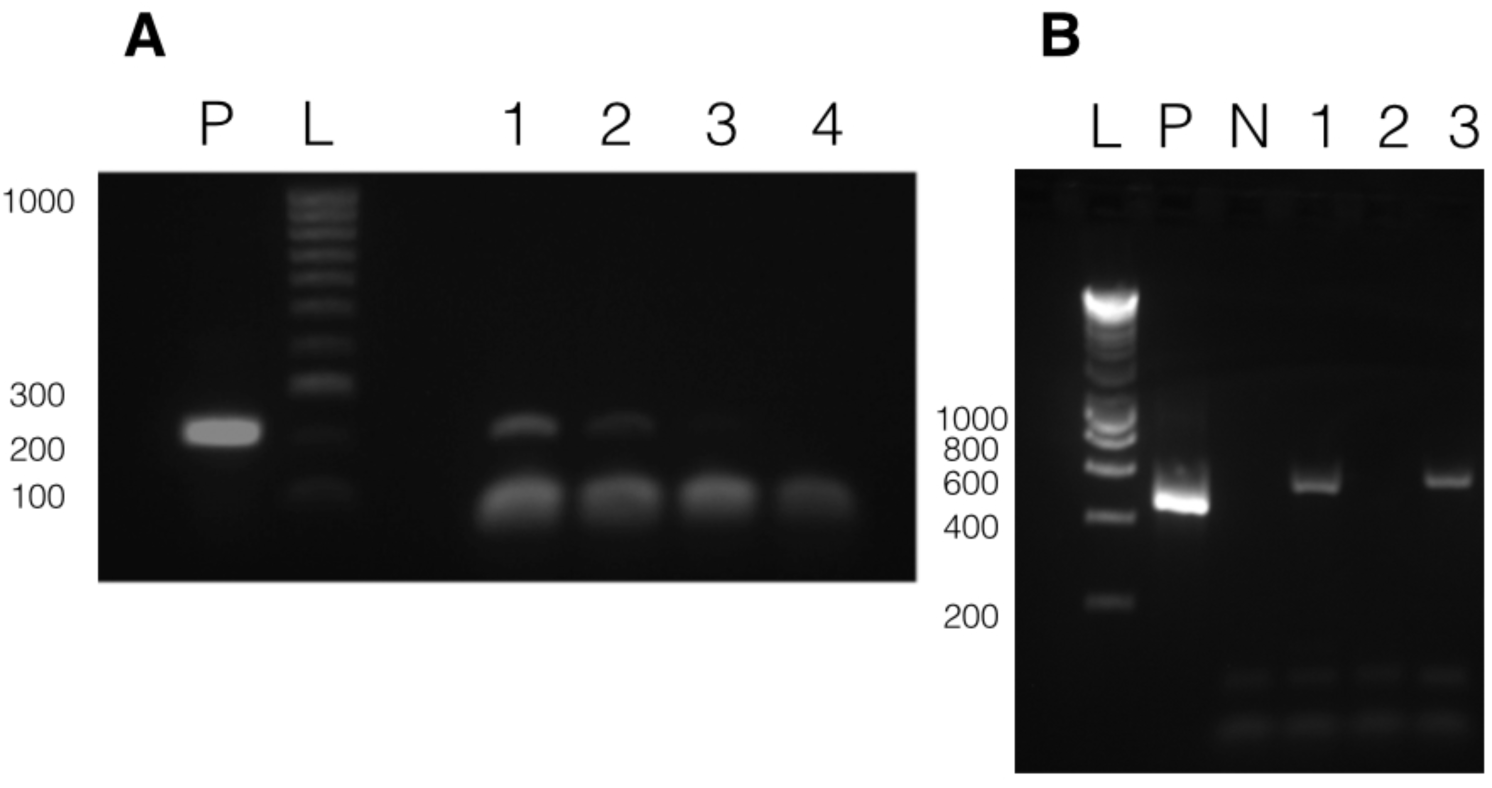
Presence of vectors in transformed *A. carterae*. Panel A shows results with pAmpPSBA. DNA was isolated from cells putatively transformed with pAmpPSBA, and a PCR reaction performed to amplify a ∼200 bp region in the plasmid. Lane P, positive control (PCR with plasmid only), Lane L, Hyperladder 100 bp (Bioline) marker, Lane 1, transformed cell line A5.1, Lane 2, transformed cell line A5.2, Lane 3, transformed cell line A5.1 but without cells being broken open prior to DNA extraction, Lane 4, wild type (i.e. untransformed cells). **Panel B shows results with pAmpChl.** DNA was isolated from cells putatively transformed with pAmpChl, and a PCR reaction performed to amplify a 580 bp region in the plasmid. Lane L, Hyperladder 1kb (Bioline) marker, Lane P, positive control (PCR with plasmid only), Lane N, negative control (no template), Lane 1, transformed cell line C1A.1, Lane 2, transformed cell line C1A.2, Lane 3, transformed cell line C1A.3. Apparent differences in mobility between bands in lanes P, 1 and 3 are due to gel loading.

To test if the pAmpChl artificial minicircle was present, DNA was isolated from three chloramphenicol-selected *A. carterae* lines (C1A.1, C1A.2 and C1A.3) after 35 days of selection. PCR using primers within the chloramphenicol resistance gene (CAT-F and CAT-R) was carried out. A positive control was included using the pAmpChl artificial minicircle. A single product was obtained from lines C1A.1 and C1A.3, with no product detected from line C1A.2 (Figure 2B). Products from lines C1A.1 and C1A.3 were cloned and sequenced. The sequence was the same as that expected from pAmpChl, which confirmed that the chloramphenicol-resistant *A. carterae* did indeed contain the pAmpChl plasmid.

### Transcription of the artificial minicircles

In order to test if transcripts from the two artificial minicircles could be detected in the putatively transformed lines, total RNA was extracted and purified. cDNA was synthesized using RNA from atrazine-selected cultures from experiment A6 (lines A6.1, A6.2, A6.3 and untransformed) and random hexamer primers, followed by PCR with the specific primers MC-pG-F-II and MC-pG-R-II. cDNA was synthesised using RNA from chloramphenicol-selected culture lines C3.1, C3.2 and C3.3 and the gene-specific primer CAT-FSS, followed by a nested PCR strategy. Primers CAT-F and CAT-R were used in the first round of PCR (30 cycles). 1 μl of PCR product was used as template for the second round of PCR (10 cycles) with primers CAT-F-Nest and CAT-R-Nest. Negative controls, which omitted the reverse transcriptase, were included for all RT-PCRs. A positive control was included using the pAmpPSBA or pAmpChl artificial minicircle.

RT-PCRs using RNA from the three atrazine-selected lines in Experiment A6 all yielded a band consistent with the size of the positive control, Figure 3A. The DNA in the bands was extracted, cloned and sequenced. The sequence of all three matched the pAmpPSBA artificial minicircle, confirming that it was transcribed. The sequence spanned the site of the atrazine resistance mutations and included the expected sequence alterations. The negative control yielded no PCR products (data not shown). The same results were obtained for three lines in each of Experiments A7 and A8 (data not shown).

RT-PCRs using RNA from the three chloramphenicol-selected lines (C3.1-C3.3) yielded bands from two of the three cell lines in Experiment C3 (Figure 3B). The PCR products were sequenced directly and shown to correspond to the pAmpChl minicircle. The negative control yielded no PCR product. Two of three lines in Experiment C2 yieled bands in RT-PCRs (data not shown).

**Figure 3.**
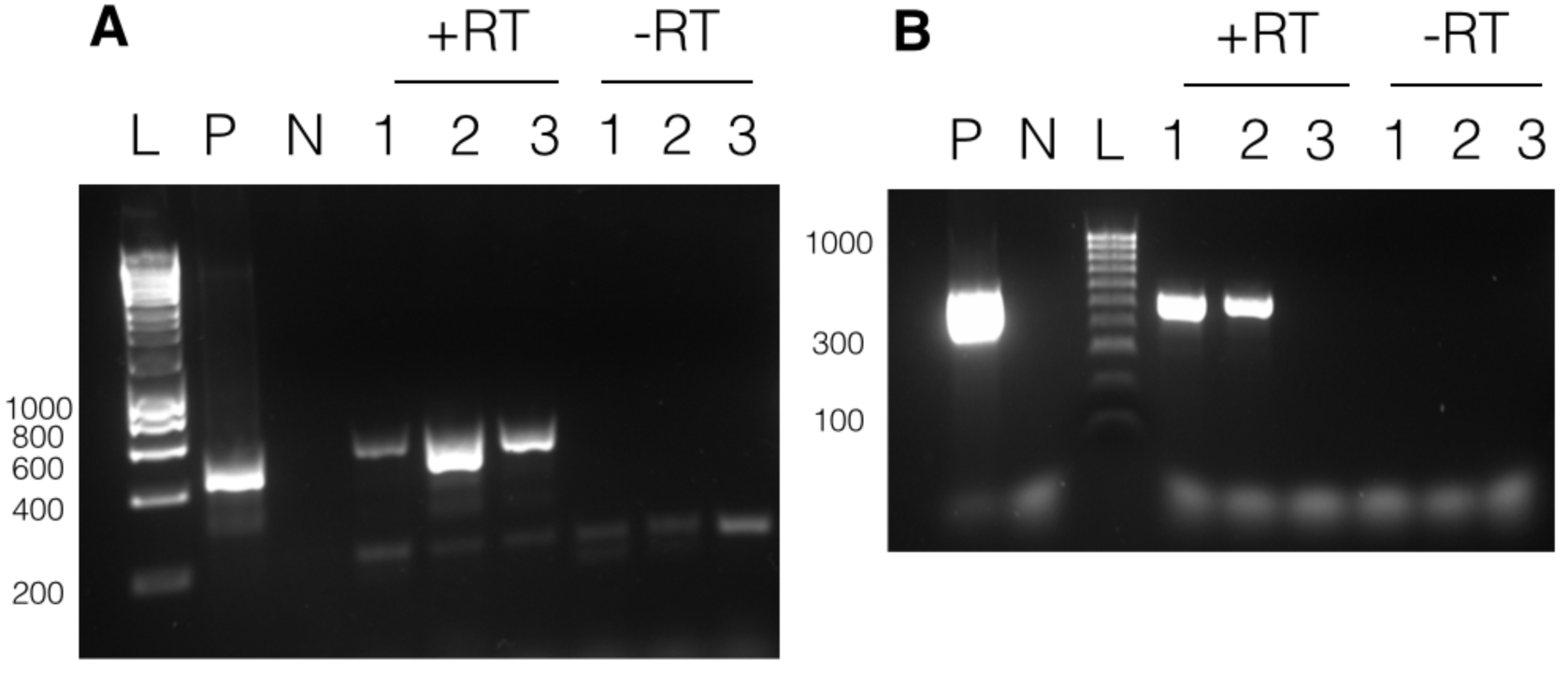
Transcription of minicircles. Panel A shows results with pAmpPSBA. RNA was extracted from cells putatively transformed with pAmpPSBA, and RT-PCR performed to amplify a ∼500 bp region. Lane L, Hyperladder 1kb (Promega), Lane P, positive control (PCR from plasmid DNA), Lane N, negative control (no template), Lanes 1-3 show products with RNA from three different pAmpPSBA-transformed cell lines (A6.1, A6.2 and A6.3) shown with reverse transcriptase (+RT) and without (-RT). **Panel B shows results with pAmpChl.** RNA was extracted from cells putatively transformed with pAMPChl, and RT-PCR performed to amplify a 580 bp region. Lane L, Hyperladder 100 bp (Bioline), Lane P, positive control (PCR from plasmid DNA), Lane N, negative control (no template), Lanes 1-3 show products with three different pAmpChl-transformed cell lines (C3.1, C3.2 and C3.3) with reverse transcriptase (+RT) or without (-RT).

### Stability of transformation

To test if the atrazine-resistance phenotype transformation of the dinoflagellate chloroplast was stable under low-level selection, cells shooting with gold particles carrying pAmpPSBA and cultured under continuous atrazine selection at 1 μg ml^−1^ (experiment A9 in Table 1). Cell counts increased over time, though at a rate much lower (∼10%) than untransformed cells under no selection. An untransformed cell line was also maintained, which survived under the same atrazine concentration (1 μg ml^−1^). Both cell lines were subcultured at eight week intervals. After three months, cells were harvested and DNA was isolated. PCR using the primers MC-pG-F and MC-pG-R was carried out (as above). A positive control PCR was included using the pAmpPSBA vector, and PCR with DNA isolated from the untransformed, wild type cells maintained at non-lethal atrazine concentration was included as a negative control (Figure 4A). A single product, of expected size, was obtained using the three transformed cell lines, with no product detected from the untransformed cells. DNA sequencing confirmed that the products from all three lines corresponded to pAmpPSBA. This showed that the transformation of *A. carterae* with pAmpPSBA was stable at a non-lethal atrazine concentration.

**Figure 4.**
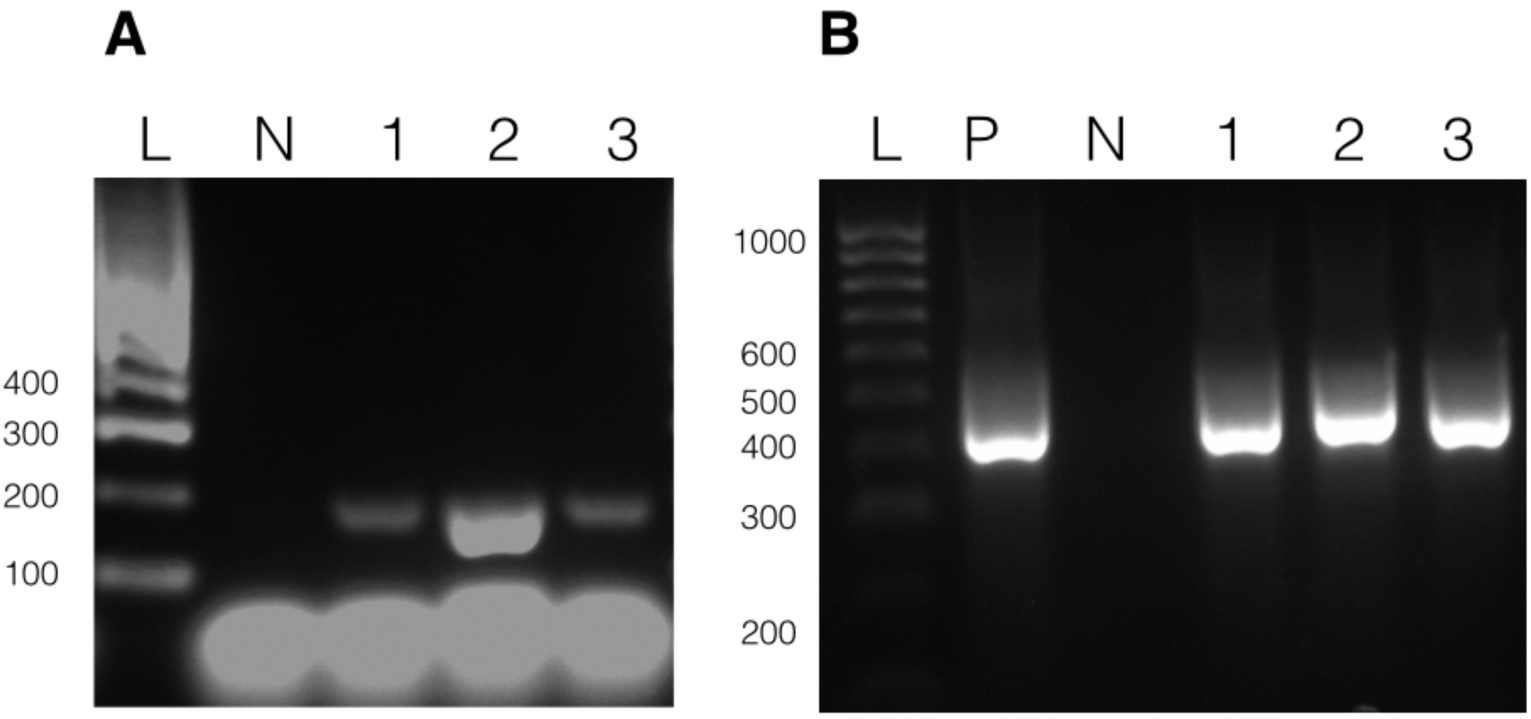
Long term stability of transformation. DNA was isolated from cells putatively transformed with pAmpPSBA or pAmpChl and maintained under selection 3 months (Experiments A9 and C4). **Panel A shows PCR to amplify a 200 bp region of pAmpPSBA.** Lane L, Hyperladder 100bp (Bioline) marker, Lane N, untransformed cells, Lane 1, transformed cell line A9.1, Lane 2, transformed cell line A9.2, Lane 3, transformed cell line A9.3. **Panel A shows PCR to amplify a 560 bp region of pAmpChl.** Lane L, Hyperladder 100bp (Bioline) marker, Lane P, positive control (pAmpChl), Lane N, untransformed cells, Lane 1, transformed cell line C4.1, Lane 2, transformed cell line C4.2, Lane 3, transformed cell line C4.3.

To test if the chloramphenicol-resistance phenotype transformation of the dinoflagellate chloroplast was stable, three lines were generated by shooting with gold particles carrying pAmpChl and cultured under continuous chloramphicol selection at 20 μg ml^-1^ (Experiment C4 in Table 2) with subculturing every 14 days. After 18 weeks, a sample of cells was harvested from each line and DNA was isolated. A PCR reaction using the primers CAT-F-Nest and CAT-R-Nest was carried out on each sample. A positive control PCR was included using the pAmpChl vector (Figure 4B). A single product, of expected size, was obtained for each of the three transformed cell lines. DNA sequencing confirmed that the products from all three lines corresponded to pAmpChl. No band was seen in a wild-type PCR carried out with the same primers on wild-type cells (data not shown), confirming that the band can only have arisen from pAmpChl. The cultures remain alive at the time of writing, 20 week post-transformation. These results show that there was stable transformation of *A. carterae* with pAmpChl.

## Discussion

Here, we present evidence for the first stable transformation of the dinoflagellate chloroplast genome. By making use of the plasmid-like fragmented chloroplast genome, we have introduced both modified versions of existing sequences and heterologous genes, and have shown that the genes are transcribed and confer the expected phenotypes. Stable transformation was achieved with two separate artificial minicircles, one encoding a modified *psbA* gene designed to confer atrazine tolerance and another encoding chloramphenicol resistance, with cultures living at least five months. In addition, *A. carterae* cells transformed with the modified *psbA* gene (atrazine tolerance) survived under non-lethal concentrations of atrazine for at least three months, retaining the modified gene, indicating that the transformation is stable even under low levels of selection.

The results suggest it is important to titrate the concentration of selective agents used. With an atrazine concentration of 2.5 μg ml^−1^ some of the transformed cultures did not survive more than a few days longer than untransformed ones, suggesting that the modified PsbA was at least partially inhibited at that higher atrazine concentration. In addition, it is possible that a background of minicircles containing wild type *psbA* genes competed with the introduced artificial minicircles for replication or transcription factors, making it difficult for adequate levels of atrazine-insensitive PsbA to be maintained to cope with the higher atrazine concentration. With chloramphenicol concentrations of 30 μg ml^−1^ or above, the survival of transformed and untransformed strains was similar. However, at 20 μg ml^−1^ or lower the transformed cultures outlasted the untransformed ones, and some were able to survive apparently indefinitely.

The ability to modify the dinoflagellate chloroplast genome will be of enormous value in many areas of dinoflagellate biology. Modification of existing minicircles should allow us to study many other aspects of this highly unusual chloroplast genome, such as the promoter regions of the genes. For example, many chloroplast genes are down-regulated under high temperature stress^21^. Little is known about how transcription is regulated, or initiated^13^, though it is assumed that initiation occurs in the conserved core region of the minicircle^15,22^. We also do not know how the minicircles are replicated^23^, though again it is assumed the core region is important. It will now be possible to mutate this core region to determine which sections are important. The ability to express heterologous proteins will be of great value in studying a wide range of other aspects of dinoflagellate chloroplast biology. The ability to express modified forms of the PsbA protein will be of particular value in studying the role of this protein in the response by dinoflagellates to the disturbances that are believed to precipitate coral bleaching.

## Acknowledgements

This research is funded by the Gordon and Betty Moore Foundation through Grant GBMF4976 to CJH, RFW, SP and MA. JEC was supported by the King Abdullah University of Science and Technology (KAUST) Office of Sponsored Research (OSR) under Award No. URF/1/2216-01-01.

